# The envelope cytoplasmic tail regulates HIV-1 assembly and spread

**DOI:** 10.1101/2021.02.08.430194

**Authors:** Xenia Snetkov, Tafhima Haider, Dejan Mesner, Nicholas Groves, Schuyler van Engelenburg, Clare Jolly

**Author notes:** Corresponding author Dr Clare Jolly Division of Infection and Immunity, University College London, Gower Street WC1E 6BT, London, United Kingdom.

## Abstract

The HIV-1 envelope (Env) is an essential determinant of viral infectivity, tropism and spread between T cells. Lentiviral Env contain an unusually long 150 amino acid cytoplasmic tail (EnvCT) but the function of the EnvCT and conserved domains within it remain largely uncharacterised. Here we identified a highly conserved tryptophan motif at position 757 (W757) in the LLP-2 alpha helix of the EnvCT as a key determinant for HIV-1 replication and spread between T cells. Strikingly we find that mutating W757 had wide-ranging consequences including altering Env mobility in the plasma membrane, preventing Env and Gag recruitment to sites of cell-cell contact for virological synapse (VS) formation and cell-cell spread, and impeding viral fusion. Notably, W757 was also required for efficient virus budding, revealing a previously unappreciated role for the EnvCT in regulating HIV-1 assembly and egress. We conclude that W757 is a key residue that stabilises the structural integrity and function of Env, consistent with the recent model that this region of the EnvCT acts as a critical supporting baseplate for Env.

## Introduction

Virus assembly is a well-orchestrated event in which all components must be temporally and spatially recruited to the right place at the right time to regulate successfully infectious viral egress and spreading infection. For the lentivirus Human Immunodeficiency Virus type-1 (HIV-1) virus, replication predominantly takes place in CD4+ T cells with virus assembly and budding occurring at the plasma membrane. HIV-1 has two major structural proteins: the envelope glycoprotein (Env) that is the viral attachment receptor that engages CD4 and co-receptor (CCR5 or CXCR4) and mediates fusion of the viral lipid membrane and the target cell plasma membrane to mediate entry; and the Gag polyprotein (comprising of capsid, matrix, nucleocapsid and p6) that forms the viral core. Env and Gag are trafficked independently to sites of lentiviral assembly at the plasma membrane. During HIV-1 assembly, trimers of Gag matrix (MA) assemble at the PM forming a lattice of hexamers which directly interact with phosphatidylinositol-4,5-bisphosphate (PIP2) in the membrane (Ono *et al*, 2004; Alfadhli *et al*, 2009). Recent super-resolution imaging has shown that the Gag lattice forms early during the course of viral assembly with Env arriving at a later stage (Buttler *et al*, 2018). Self-multimerisation of Gag, lattice associated Env, and the host cell ESCRT machinery together appear to drives virus assembly and budding (Stuchell *et al*, 2004; Morita *et al*, 2011; Prescher *et al*, 2015; Van Engelenburg *et al*, 2014). Whether the EnvCT interacts with Gag MA, how exactly Env gets incorporated into virions and whether Env plays any role in regulating virus assembly remains unclear (Checkley *et al*, 2011; Murphy & Saad, 2020).

Env is composed of gp120 domain that mediates receptor binding, and the gp41 domain that is composed of an extracellular and transmembrane domain, and a long cytoplasmic tail of 150 amino acids. Conservation of the long lentiviral envelope cytoplasmic tail (EnvCT) suggests the presence of key determinants within the CT that are essential for efficient lentiviral replication and spread (Murakami & Freed, 2000). Indeed, the importance of the EnvCT is best demonstrated by the observation that when the C-terminal EnvCT is truncated (CTΔ144 virus), HIV-1 is unable to spread in T cell lines and primary CD4 T cells due to an Env incorporation defect (Murakami & Freed, 2000). The N terminus of the EnvCT consists of short membrane proximal unstructured loop containing a YxxL endocytic motif that binds the clathrin adaptor, AP-2 (Berlioz-Torrent *et al*, 1999; Ohno *et al*, 1997). Rapid endocytosis of Env from the plasma membrane via the YxxL and an additional C-terminal dileucine motif limits the amount of Env present of the surface of infected cells (Berlioz-Torrent *et al*, 1999; Ohno *et al*, 1997; Byland *et al*, 2007; Wyss *et al*, 2001; Bhakta *et al*, 2011). This is thought to protect infected cells from detection by the immune system and limit the amount of Env packaged into nascent virions to aid immune evasion (Stano *et al*, 2017; Rusert *et al*, 2016). Following endocytosis from the plasma membrane, retrograde and recycling trafficking pathways are proposed to coordinate Env recycling back to the cell surface (Groppelli *et al*, 2014; Qi *et al*, 2013; Wang *et al*, 2020; Bhakta *et al*, 2011; Byland *et al*, 2007). Previous work has shown that endocytosis mediated by Y712 also helps direct virus budding to the basolateral membrane in epithelial cells and also influences the site of virus budding in T cells, presumably by allowing endocytosed Env to access intracellular trafficking machinery for correct sorting back to virus assembly sites (Deschambeault *et al*, 1999; Groppelli *et al*, 2014; Kirschman *et al*, 2017). Thus, while the biology of the EnvCT remains to be fully-elucidated, the absolute dependence on the EnvCT for infectious virus assembly in T cells argues against a passive incorporation model and highlights the importance of the EnvCT in HIV-1 replication and spread. Notably, the first structural data of the EnvCT revealed that the majority of the long EnvCT resides buried in the membrane (Murphy *et al*, 2017). This is intriguing given that the EnvCT of related retroviruses is considerably shorter at 30-40 amino acids (Postler & Desrosiers, 2013). Thus, it is likely that the HIV-1 EnvCT contains additional undefined motifs that regulate Env trafficking, localisation to virus assembly sites and virion incorporation.

HIV-1 spreads between CD4+ T cells via two distinct mechanisms; either by cell-free infection or by direct cell-to-cell transmission (reviewed in Sattentau, 2008). The latter is the dominant mode of HIV-1 spread (Jolly *et al*, 2007a, 2004; Chen *et al*, 2007; Sourisseau *et al*, 2007) in which the virus exploits the tight packing of CD4+ T cells in lymphoid organs and their propensity for frequent interactions to rapidly spread from one cell to another (Murooka *et al*, 2012). This occurs following physical contact between an HIV-1 infected donor T cells and the uninfected target T cells and is termed the virological synapse (VS) (Jolly *et al*, 2004). VS formation is characterised by the recruitment of HIV-1 Env and Gag to sites of cell-cell contact leading to polarised virus assembly and budding that is focused towards the target cell (Jolly *et al*, 2004; Hübner *et al*, 2009; Chen *et al*, 2007; Rudnicka *et al*, 2009). Thus, cell-cell spread is an active and regulated process that is triggered by cell-cell contact, but the mechanisms by which Env and Gag are directed to the VS and how virus assembly is orchestrated during cell-cell spread remains to be fully-elucidated. Understanding this is important, not only because may it reveal new therapeutic options to limit HIV-1 spread by targeting the late-stages of HIV-1 replication, but it also provides a tractable and valuable experimental model to interrogate how HIV-1 assembly is regulated and reveal novel viral determinants.

Here we have sought to provide molecular insight into the processes of HIV-1 assembly and spread, specifically the role of the enigmatic EnvCT. To this end, we have identified a highly conserved tryptophan residue at position 757 within the first helix of the EnvCT (LLP2) that mediates HIV-1 spread between T cells, regulating Env and Gag recruitment to virus assembly sites, limiting lateral diffusion of Env within the plasma membrane and regulating Env fusogenecity. Thus our data identify a highly conserved site within the EnvCT that plays a key role in regulating multiple Env functions, and notably reveal a new and unappreciated role for the EnvCT in triggering HIV-1 assembly in T cells.

## Results

### A conserved tryptophan residue in the EnvCT is required for HIV-1 cell-cell spread and virological synapse formation

To identify novel determinants in EnvCT that play a role in HIV-1 replication and spread we focused on tryptophan (W) residues that possess a large bulky hydrophobic side chain and act as key mediators of protein-protein and protein-lipid interactions, including the within the W-rich HIV-1 MPER (Salzwedel *et al*, 1999; Muñoz-Barroso *et al*, 1999). The 150 amino acid HIV-1 EnvCT contains five tryptophan residues at positions 757, 790, 796, 797 and 803 (Fig. 1A). Analysis of 5916 HIV-1 Env sequences from the Los Alamos database (http://www.hiv.lanl.gov/) revealed that three of these (W757, W790, and W803) are highly-conserved amongst HIV-1 isolates (Fig. 1A) (Crooks *et al*, 2004; Foley *et al*, 2018). We focused on the two tryptophan residues at W757 and W790, given that disruption of Y802-W803 has been previously characterised and shown to reduce Env incorporation and infectivity (Lambelé *et al*, 2007; Blot *et al*, 2003). Using a mutagenesis approach, the first tryptophan at the beginning of the LLP-2 helix of the cytoplasmic tail of HIV-1 envelope (W757) was substituted for alanine in full length replication competent HIV-1 NL4.3 and the effects on viral replication and cell-cell spread were examined. To normalise infection (and obviate any differences in the proportion of HIV-1 infected cells that may bias analysis), HIV-1 wild type (WT) and the tryptophan mutant (W757A) were pseudotyped with VSV-G to achieve equivalent initial infection of T cells. We focused on cell-cell spread that is the dominant mode of HIV-1 dissemination between T cells and takes place at the VS. Jurkat cells (donors) were infected with WT or W757A virus for up to 48 hours, mixed with dye-labelled uninfected Jurkat cells (targets) and cell-cell interactions and VS formation were quantified. Consistent with previous reports, co-culturing HIV-1 WT infected donors and uninfected targets resulted in 62% of cell-cell contacts forming a VS, evidenced by polarisation of Env and Gag on infected cells to the sites of contact with a target cell (Fig.1B, C). Strikingly, substituting the tryptophan at position 757, with an alanine (W757A virus) abolished VS formation resulting in loss of both Env and Gag recruitment to the contact zone (Fig. 1B, C). Importantly, the frequency of cell-cell contacts formed between donor and target T cells remained unchanged, implicating a specific defect in VS formation and not an inability of cells to interact (Fig. 1D). By contrast to WT Env, W757A Env displayed a more punctate, uninform distribution around the plasma membrane that was not enriched at the contact zone (Fig. 1B, E). As expected, the inability of the HIV-1 Env W757A mutant to form VS resulted in a significant defect in HIV-1 cell-cell spread. Co-culture of WT infected Jurkat T cells and uninfected target T cells resulted in 55% of target T cells becoming infected after 24h, by contrast to W757A virus that was unable to spread (Fig. 1F and Supplementary Fig. 1). Similar results were obtained when spreading infection was performed using primary CD4+ T cells; however, in this case the defect in W757A virus was even more pronounced, with no increase in viral spread to target cells evident up to 50hr post-infection (Fig. 1F and Supplementary Fig. 1). By contrast to the W757A mutant, the W790A mutant displayed no defect in VS formation or cell-cell spread (Supplementary Fig. 2). Therefore, we focused on understanding the specific requirement for W757 on Env biology and viral spread.

**Figure 1.**
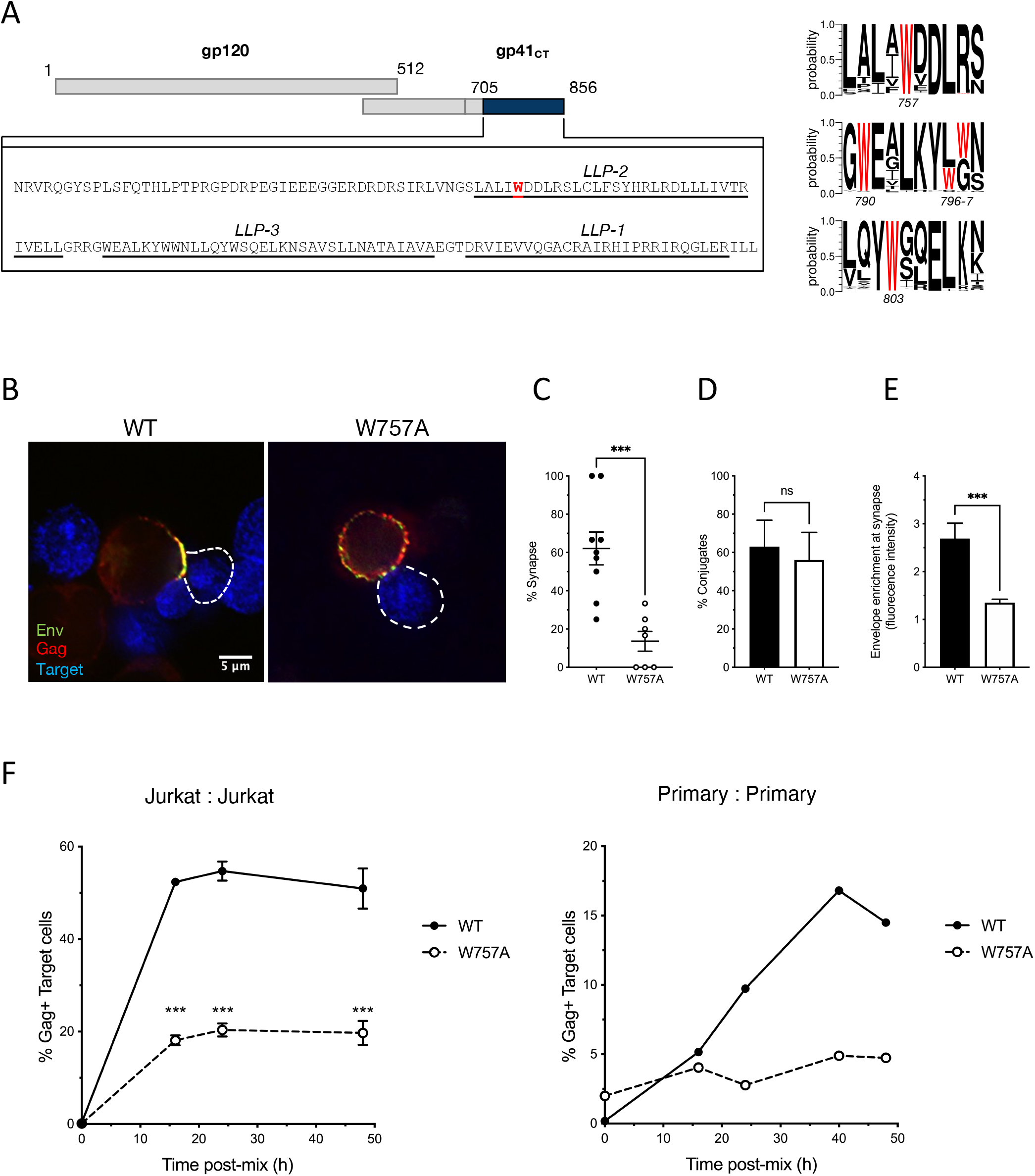
EnvCT W757A does not polarise to viral synapses and leads to a defect in cell-cell spread. (A) Schematic showing the HIV-1 NL4.3 EnvCT sequence and location of W757 in the LLP2 alpha helix. Logo plots (generated using WebLogo3) indicate the probability of 5916 HIV-1 Env sequences analysed in the LANL database containing a W residue at the indicated positions. **(B)** Synapse formation between Jurkat cells infected with WT NL4.3 (left panel) or the W757A mutant (right panel) and uninfected target cells. NL4-3 virus pseudotyped with VSVg to normalise viral entry was used to infect donor Jurkat cells. Representative image is an *xy* slice through the middle of a cell-cell contact between an infected donor cell (Env, green; Gag, red) and a target cell stained with a cell trace dye (blue). **(C)** Percent of contacts that exhibited synapse formation (defined by polarisation of both Env and Gag to the contact site). **(D)** Quantification of the percentage of infected donor cells (Gag+) in contact with target cells (blue). Data show the mean and SEM from n=54-69 infected donor cells compared using a two-tailed paired t test (ns, not significant; *** p < 0.001). **(E)** Signal intensity enrichment at the synapse, relative to a synapse distal site was determined from 20 images using ImageJ analysis. **(F)** Infected donor cells were mixed in a 1:1 ratio with target cells (stained with cell trace far red) and viral spread was measured by intracellular Gag staining in target cells by flow cytometry. Data show the percent of Gag+ target cells over time. Jurkat cells (left) and primary CD4+ cells (right). Data show the mean and SEM from four independent Jurkat cells experiments analysed using multiple paired t tests (ns, not significant; *** p < 0.001). For primary CD4^+^ T cells a representative donor from three unique donors is shown.

### Single molecule tracking of Env at virus assembly sites

To explain the defect in spreading infection of W757A virus, and the inability of Env to polarise to virus assembly sites at the VS, single particle tracking (SPT) was performed on Env trimers proximal and distal to super-resolved sites of virus assembly to interrogate the mobility of wild-type EnvCT and W757A mutants (Groves *et al*, 2020). This approach has previously elucidated the dynamics of Env mobility at the cell surface, revealing that during nascent virus assembly Env becomes trapped or corralled in the Gag lattice to facilitate incorporation into virions (Pezeshkian *et al*, 2019; Buttler *et al*, 2018) (Fig. 2A). Importantly, the lateral diffusion of Env within and outside virus assembly sites was shown to be dependent on the EnvCT, as well as Gag MA (Pezeshkian *et al*, 2019). Thus, we hypothesised that failure of Env W757A to localise to virus assembly sites and polarise to the VS may be explained by a lattice trapping defect, leading to inefficient incorporation of Env into the budding Gag lattice. Exploring this notion, single-molecule tracking of Env and super-resolution imaging Gag assembly sites revealed that trimers of Env W757A were significantly more mobile in the PM compared to WT Env, with Env W757A diffusing more freely when distal to the Gag lattice (lower panel -Fig. 2A, B; D_apparent_ = 0.07 ± 0.17 for Env WT versus D_apparent_ = 0.05 ± 0.10 for Env W757, *P* < 0.0001; S_MSS_ = 0.12 ± 0.21 for Env WT versus S_MSS_ = 0.13 ± 0.22 for Env W757, *P* = 0.0017).

**Figure 2.**
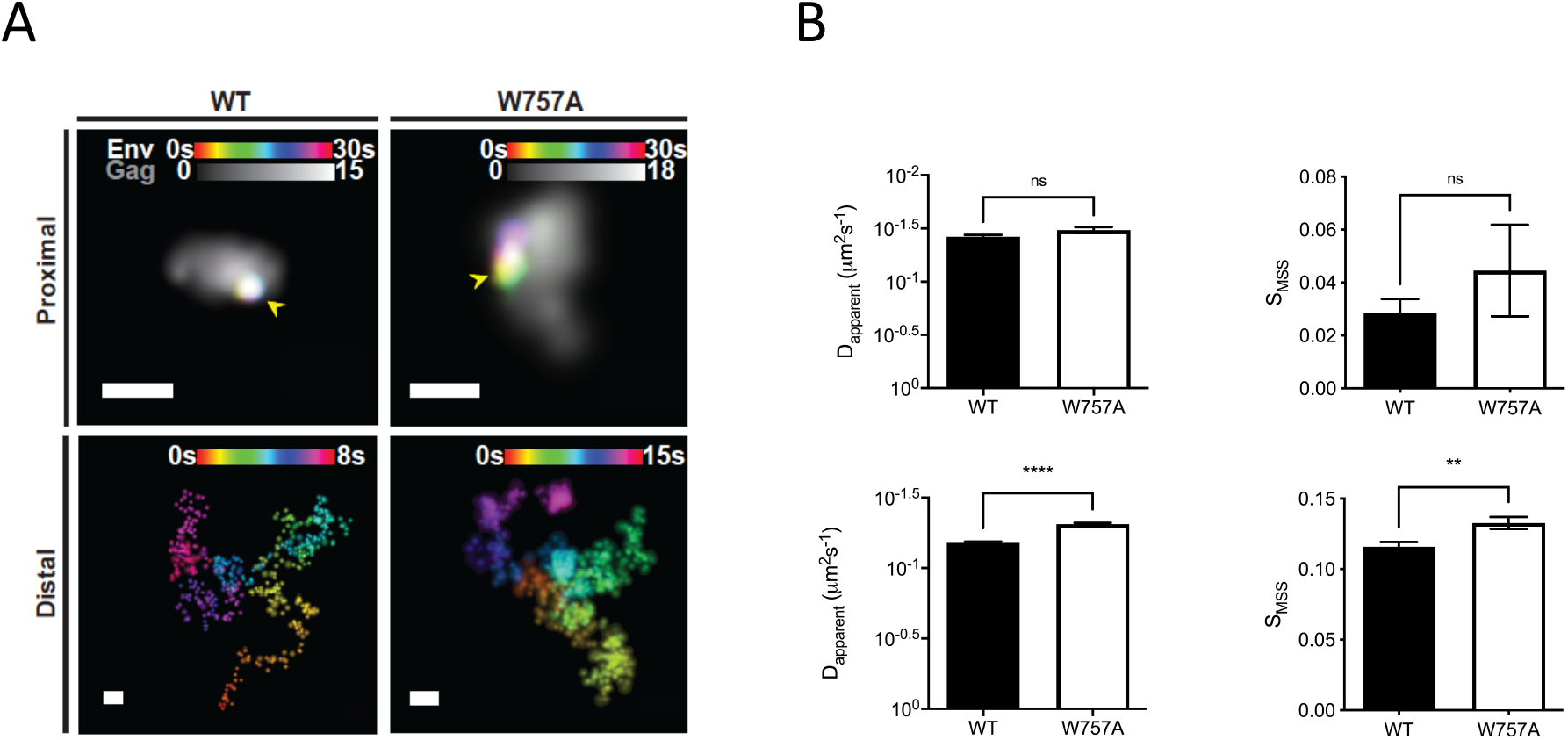
W757A EnvCT does not display a Gag lattice trapping defect at virus assembly sites. CEM-A adherent T cells were transduced with WT and W757A mutant Envelope virus and imaged after 48 hours. **(A)** Super-resolution reconstructions of Gag and lattice proximal/distal single-molecule localisations of Env are demonstrated (Scale bars = 200nm. Arrows highlight highly confined single molecule localisations of Env). Proximal and distal single-molecule localisations show the confined and freely-diffusing nature of single Env trimers, respectively. **(B)**. Wild-type and mutant Env trimers do not differ in mobility when lattice proximal (top panels) (WT: *D*_*apparent*_ = 0.038 μm^2^s^−1^, *S*_*MSS*_ = 0.01; W757A: *D*_*apparent*_ =0.033*μ*^2^*s*^−1^, *S*_*MSS*_=0.045), whereas lattice-distal diffusion differs significantly between viruses (bottom panels) (WT: *D*_*apparent*_ = 0.066 *μ*^2^*s*^−1^, *S*_*MSS*_ = 0.116; W757A: *D*_*apparent*_ = 0.049 *μ*^2^*s*^−1^, *S*_*MSS*_ = 0.133). Statistical significance was determined by unpaired two-sample t-test. 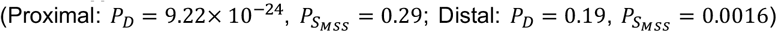.

By contrast, proximal W757A mutant trimers exhibited similar diffusion characteristics as lattice proximal WT Env trimers (D_apparent_ = 0.038±0.06 for Env WT D_apparent_ = 0.033 ± 0.03 for Env W757, *P* = 0.0693; S_MSS_ = 0.03 ± 0.13 for Env WT versus S_MSS_ = 0.05 ± 0.16 for Env W757, *P* = 0.3743), suggesting that W757A mutant trimers are not defective for lattice trapping and confinement (upper panel -Fig. 2B, C). Taken together these data suggest a role for W757 in the EnvCT in modulating Env movement within the PM.

### W757 is required for efficient HIV-1 budding and infectivity

Integrating the striking defect in cell-cell spread and VS formation with super-resolution single-molecule imaging of Env suggested that mutating W757 in the EnvCT perturbed Env mobility in the PM and therefore the temporal and spatial recruitment of Env to virus assembly sites; however, we were struck by the observation that Gag also failed to polarise to the VS when cells were infected with the W757A Env mutant virus. This suggests a hitherto unappreciated role for the EnvCT (specifically W757) in regulating Gag-dependent HIV-1 assembly and budding from T cells for subsequent viral spread. To explore this further, we quantified the cell-free budding and infectivity of virus produced by Jurkat T cells and primary CD4+ T cells infected with WT or W757A virus. Strikingly, infection of both Jurkat and primary CD4^+^ T cells with W757A virus resulted in a significant 4-5 fold reduction in HIV-1 budding compared to WT virus, quantified by SG-PERT assay for reverse transcriptase activity in culture supernatants (Fig. 3A). Importantly, mutating W757 did not result in amino acid changes to *Rev* that have been reported to mediate budding defects when the EnvCT is truncated (Durham & Chen, 2015). Measuring virus infectivity by titrating virus-containing supernatants onto HeLa TZM-bl cells and normalising the relative light units (RLU) to RT units showed that not only were fewer particles released from infected T cells, but that these viral particles were significantly less infectious (Fig. 3B). However, examining Env in virions by immunoblotting showed that incorporation of Env gp120 into viral particles was similar between WT and W757A virus (Fig. 3C) with no significant difference between these viruses in the gp120:p24 ratio in virions (Fig. 3D). Consistent with no observable Env incorporation defect, mutation of Q62 in the Gag MA, which has been shown to rescue Env incorporation defects, did not revert the W757A mutant phenotype back to WT (Tedbury *et al*, 2013; Ono *et al*, 1997)(Supplementary Fig. 3). The lack of an Env incorporation defect is also consistent with the single-molecule tracking data showing that both WT Env and W757A Env get trapped in the Gag lattice at the PM, and thus would be expected to get incorporated into budding virions (Pezeshkian *et al*, 2019). Taken together, these data suggest that W757 in the EnvCT is not only a determinant of Env recruitment to virus assembly sites, but that it also contributes to regulation of HIV-1 budding and infectivity.

**Figure 3.**
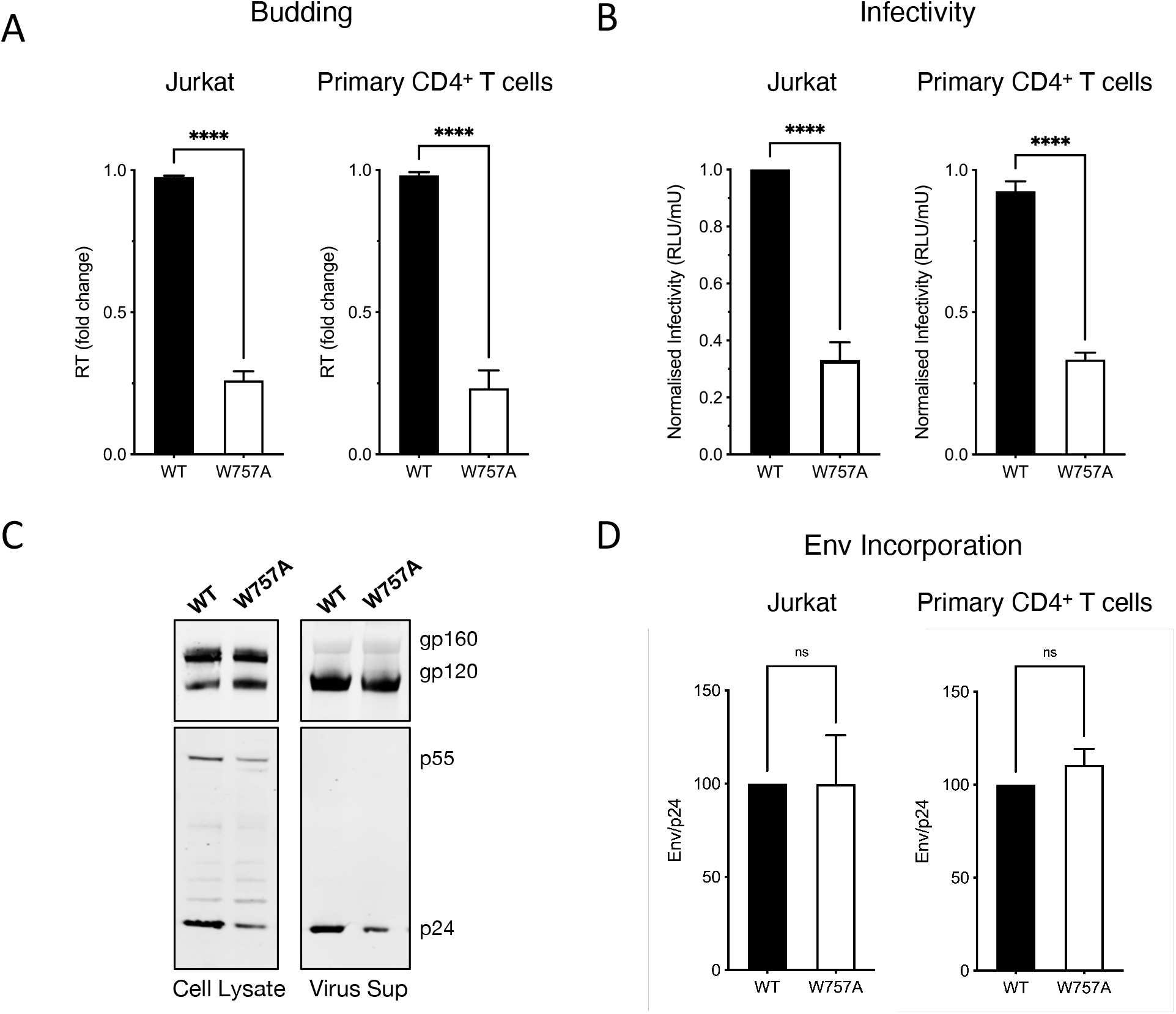
EnvCT W757 is required for HIV-1 budding and infectivity. (A) Virus release from infected Jurkat and primary CD4^+^ T cells at 48hpi measured by quantifying RT activity in viral supernatants by SG-PERT qPCR. **(B)** Infectivity of virus particles released 48hpi was measured by titrating virus on HeLa TZM-bl reporter cells and RLU normalised per RT unit to calculate relative particle infectivity. **(C)** Western blot of cell lysate and purified virus from infected primary CD4^+^ T cells at 48hpi probed with anti-Env (gp120/gp160) and anti-Gag (p55/p24). **(D)** Quantification of Env (gp120) incorporation into virions normalised to Gag (p24) density, produced in Jurkat and primary CD4^+^ T cells. Data show the mean and SEM from at least three independent experiments compared using a two-tailed paired t test (ns, not significant; **** p < 0.0001).

### Synthesis, processing, and recycling of EnvCT W757A is equivalent to that of WT EnvCT

To address the observation that W757 in the EnvCT is required for viral infectivity, but apparently not virion incorporation, we tested Env expression, processing, localisation and trafficking in infected T cells. VS formation (and subsequent cell-cell spread) is dependent on expression of the Env glycoprotein on the surface of infected cells interacting with cell surface receptors on target cells (Jolly *et al*, 2004). Immunofluorescence microscopy showed no difference in intracellular staining of WT and W757A Env (Fig. 4A), with both displaying perinuclear localisation consistent with the bulk of Env residing in Golgi compartments as expected (Blot *et al*, 2003). Immunoblotting of infected cell lysates confirmed similar levels of unprocessed Env gp160 and the furin-cleavage products gp120 and gp41 (Fig. 4B). To compare surface expression of Env on infected T cells flow cytometry analysis was performed. WT and W757A infected T cells (Jurkat and primary CD4^+^ T cells) expressed identical levels of Env on the surface of infected cells (Fig. 4C). Once at the PM, a highly conserved membrane proximal AP-2 binding motif (YxxL) acts to limit the amount of Env presented at the surface of infected cells with internalised Env being recycled back to the PM (Mesner *et al*, 2020; Groppelli *et al*, 2014; Kirschman *et al*, 2017). To test the recycling kinetics of WT and W757A Env we used a dual-fluorophore flow cytometry-based assay (Anand *et al*, 2019; Mesner *et al*, 2020). Two Env antibodies, 2G12 and PGT151 were allowed to bind surface Env at 4^°^C and detected using a PE-conjugated secondary antibody. Recycling over time was allowed to proceed at 37^°^C and the population of Env that returned to the surface was detected with a second Cy5-secondary resulting in double PE/Cy5 labelled Env. No significant difference in recycling of endocytosed Env back to the PM was seen between WT and W757A Env (Fig. 4D). Finally, we tested whether Env W757A was competent for CD4 and co-receptor (CXCR4) binding and trimer formation using neutralisation assays (Montefiori, 2009). WT and W757A viruses were both neutralised by bNAb PGT151 (that binds to the gp120/gp41 trimer interface and detects functional Env trimers (Blattner *et al*, 2014; Falkowska *et al*, 2014)), soluble CD4 (sCD4) and the CXCR4 co-receptor inhibitor AMD3100 (Donzella *et al*, 1998) (Fig. 5A). While subtle differences in neutralisation were observed between WT and W757A virus, these are insufficient to explain the striking defect in viral infectivity. In order to investigate whether a fusion defect may be responsible for the decrease in Env W757A infectivity, neutralisation with the fusion inhibitor Enfuvirtide (T20) was carried out. Strikingly Env W757A was unable to be completely neutralised with T20. Compared to WT Env, a maximum inhibition of 50% was achieved (Fig. 5A). To test whether W757 was fusion-competent the BlaM-Vpr fusion assay was used to measure the capacity of Env W757A virions to fuse with target cells, resulting in BlaM-Vpr release into the target cell and cleavage of CCF2 (Cavrois *et al*, 2002). Fig. 5B shows that Env W757A was significantly less able to mediate virion fusion compared to WT virus, demonstrating that the infectivity defect of W757A can be explained by a defect in fusion of the viral and host cell membranes and thus reduced virus entry (Fig. 5B).

**Figure 4.**
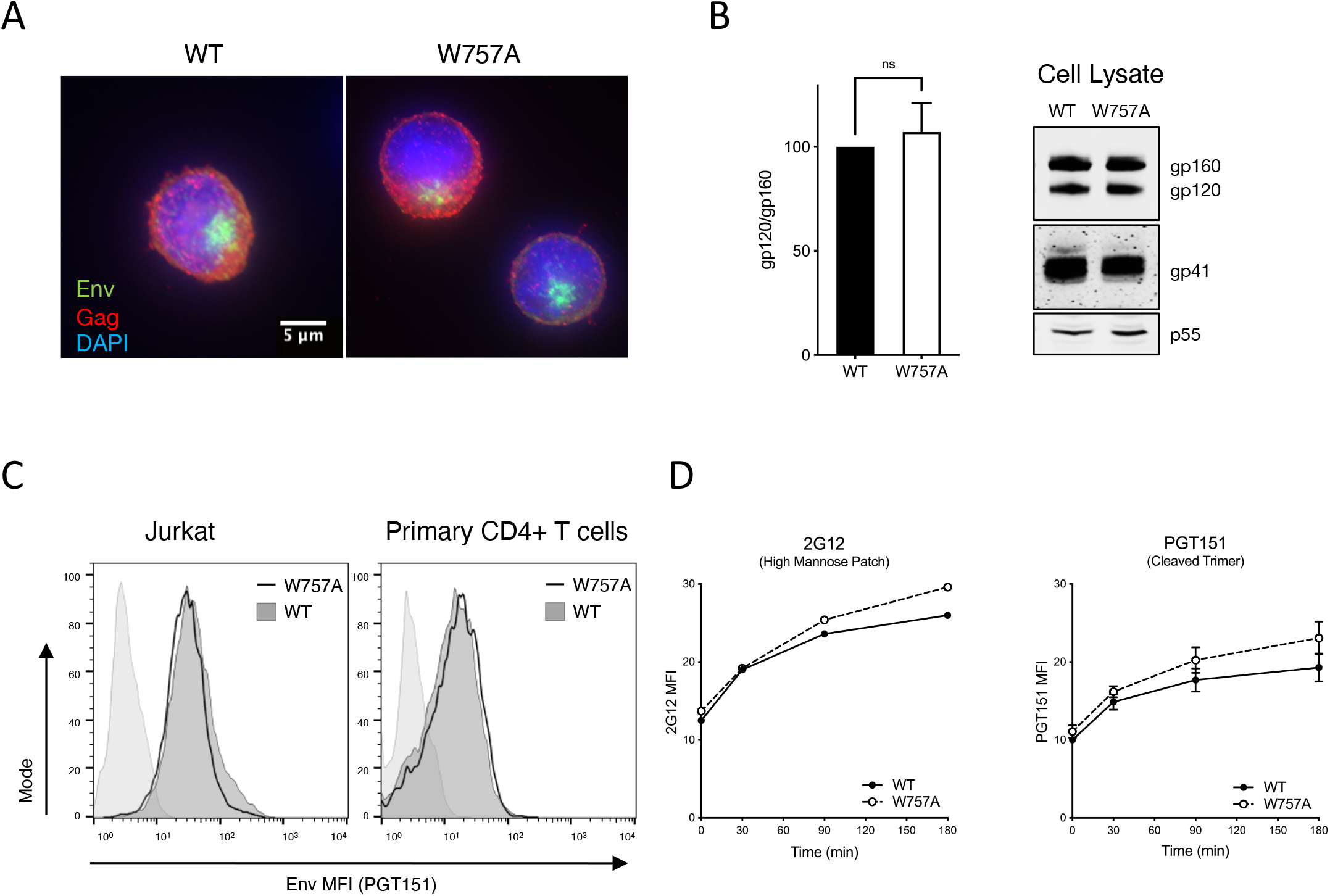
Synthesis, processing, and recycling of EnvCT W757A is equivalent to WT EnvCT. (A) Intracellular staining of Env (green) and Gag (red) in Jurkat cells infected with WT and W757A at 48hpi. **(B)** Western blot of infected Jurkat cell lysates harvested 48hpi and probed with anti-gp120, anti-gp41, and anti-Gag p55/p24. Quantification of Env processing (gp120/gp160). Data show the mean and SEM from at least three independent experiments compared using a two-tailed paired t test (ns, not significant). **(C)** Flow cytometry analysis of cell surface Env levels on cells infected with WT (black) and W757A (grey) virus measured using bNAb PGT151. Uninfected control is shown in light grey. **(D)** Time course of endocytosed Env recycling back to the plasma membrane measured by flow cytometry. Env was stained with either 2G12 or PGT151. Envelope recycling was measured by staining with envelope antibody and a PE-secondary on ice followed by Cy5 at 37°C. Recycling is shown as the MFI of PE^+^CY5^+^ signal on the surface over time. Data show the mean and SEM from three independent experiments.

**Figure 5.**
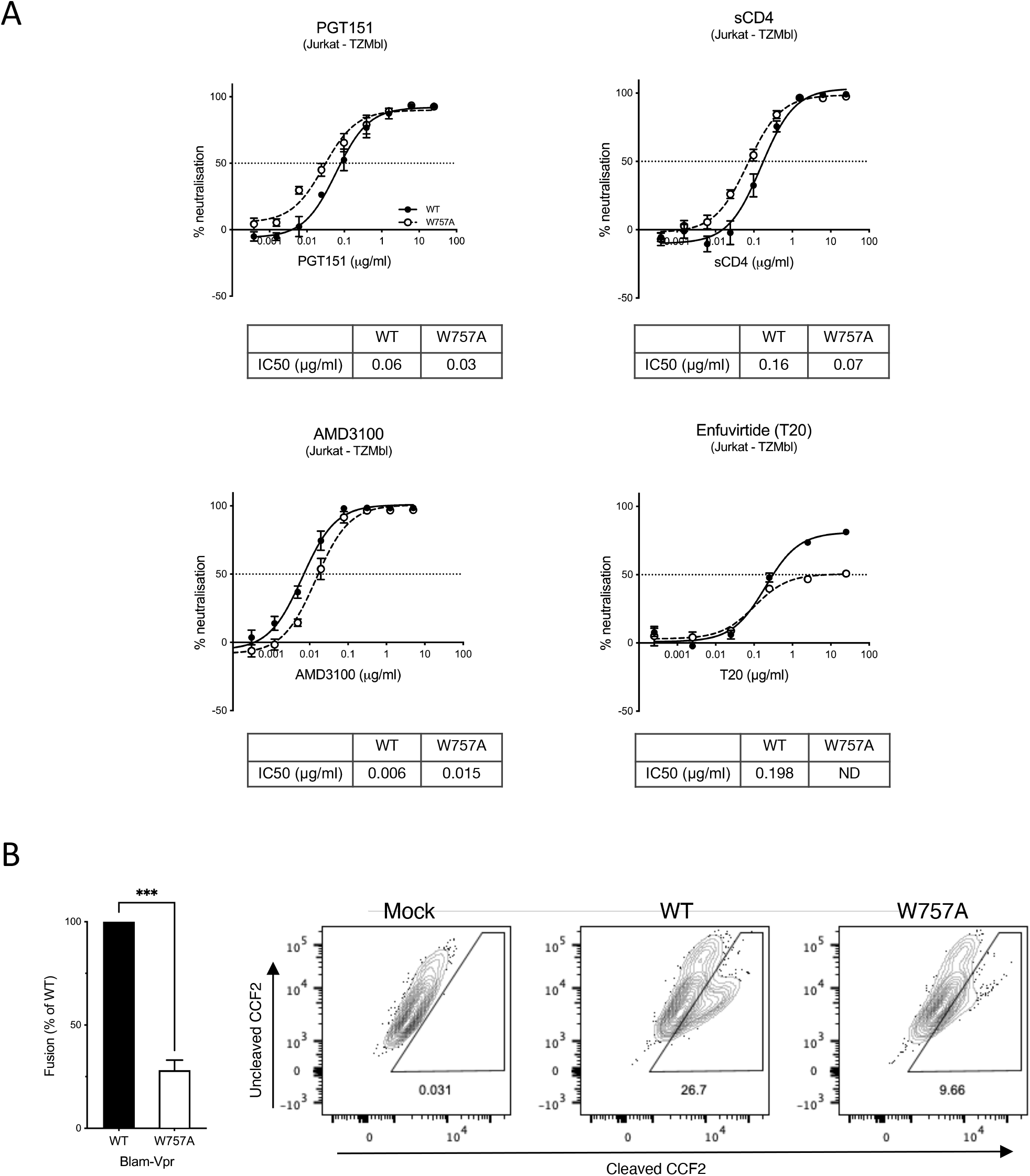
W757A virus shows a defect in viral fusion. (A) Neutralisation of WT and W757A virus (produced from Jurkat T cells) by PGT151 bNAb, sCD4, AMD3100 and T20. IC_50_ values calculated by non-linear regression analysis of the neutralisation curves. ND denotes not determined. **(B)** W757A Env is defective in viral fusion compared to WT virus as measured by the BlaM-Vpr fusion assay. Data are the mean and SEM compared using an unpaired t test (*** p< 0.001). Mock refers to co-culture between uninfected T cells and target T cells.

### Conservation of W757 across different HIV lineages and ancestors

Finally, we sought to explore conservation of W757 across HIV-1 groups and related lentiviruses. Examining 194 Env sequences at position 757 across HIV-1 group M, N, O, P, SIVgor and SIVcpz using Chromaclade (Monit *et al*, 2019; Foley *et al*, 2018) illustrated that a W at position 757 is highly conserved amongst HIV-1 group M, SIVcpz and HIV-1 group N; but not group O, or HIV-2 and the related SIVsmm and SIVmac viruses (Fig. 6A and Supplementary Fig. 4). Noting that all HIV-1 sequences (M, N, O and P) contained an aromatic residue at 757 (W or Y) we hypothesised that conservation of an aromatic residue at this position was important and that substituting the A for a Y or F would rescue virus replication and spread. Indeed, substituting the A at position 757 with a Y (HIV-1 O group) or F (another aromatic) rescued the W757 mutant and phenocopied the WT W757 virus, restoring virus budding (Fig. 6B), infectivity (Fig. 6C) and cell-cell spread (Fig. 6D). Taken together this data implicate evolutionary conservation of an aromatic residue in the EnvCT at position 757 as a potentially important feature of HIV-1.

**Figure 6.**
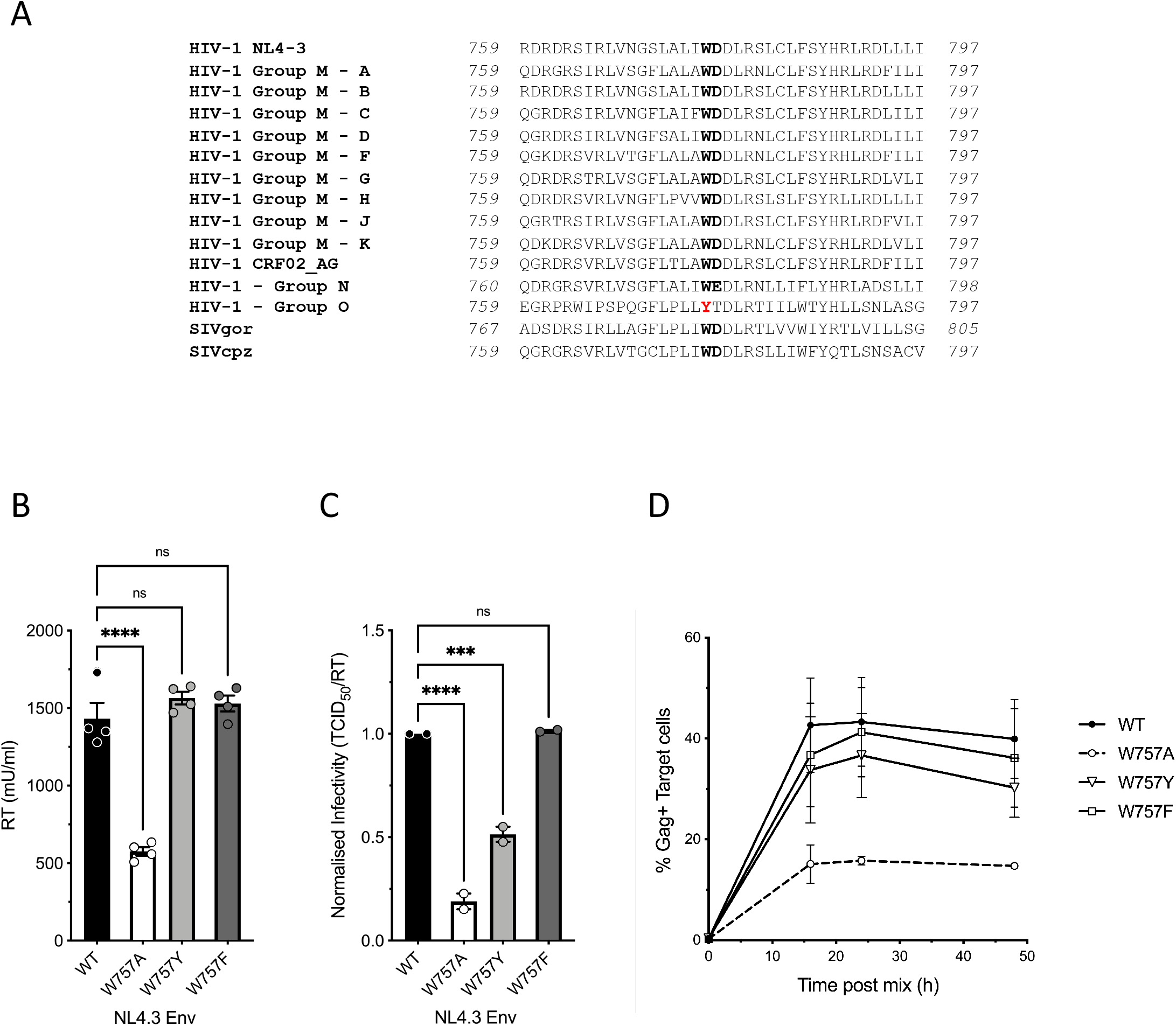
Conservation of W757 across HIV and SIV lineages. (A) Amino acid alignment of the LLP2 domain of the EnvCT from different HIV and SIV consensus sequences. See also Supplementary Figure S4. **(B-D**) Viral budding, infectivity, and cell-cell spread can be rescued by substitution at position 757 with either tyrosine Y, or phenylalanine, F. Data are the mean and SEM compared using a two-tailed paired t test (ns, not significant; *** p< 0.001).

## Discussion

Here we have uncovered a key role for a highly-conserved tryptophan residue (W) in the HIV-1 EnvCT that is essential for correct temporal and spatial regulation of HIV-1 assembly following cell-cell contact and subsequent viral spread between T cells. The mechanism by which HIV-1 assembly is regulated, and particularly the spatiotemporal events governing Env and Gag recruitment to the contact zone during synapse formation leading to polarised viral assembly remain largely unclear. Moreover, why lentiviruses conserve such an unusually long EnvCT and what role it plays on HIV-1 assembly and infectivity remains enigmatic. Given the dominance of highly-efficient cell-cell spread in viral dissemination, understanding how this process is coordinated is important (Jolly *et al*, 2004; Chen *et al*, 2007; Sourisseau *et al*, 2007). Multiple (but not mutually exclusive) mechanisms for Env and Gag recruitment to sites of virus assembly at the VS have been proposed. In the case of Env, there is evidence for Env being directed to the VS via lateral diffusion, outward transport from polarised secretory apparatus and recycling (Wang *et al*, 2020; Kirschman *et al*, 2017; Starling *et al*, 2016). By contrast, Gag is thought to be recruited independently of Env with evidence for lateral diffusion (Hübner *et al*, 2009) and polarised trafficking (Groppelli *et al*, 2015) to the contact zone. By coupling viral assays with single particle tracking at superresolved virus assembly sites we find that the EnvCT via W757 is required to direct Env and Gag to sites of cell-cell contact and mediate VS formation and spreading infection in T cells. Notably, single-molecule tracking revealed that mutating W757 resulted in Env diffusing more freely within the PM compared to WT Env. Structural insights of the EnvCT by solution state NMR revealed that much of the EnvCT associates closely with cellular membranes, and W757 at the start of the first alpha helix (LLP-2) in the EnvCT appears to provide a crucial anchor point in order to regulate membrane diffusion (Supplementary Fig. 5) (Murphy *et al*, 2017; Piai *et al*, 2020). Perturbation of this W757 anchor point may lead to changes in the quaternary structure of the putative LLP-2 baseplate and could explain an increase in PM diffusivity of the W757A mutant Env trimer compared to the native EnvCT (Piai *et al*, 2020). Disruption of the EnvCT baseplate by W757A may not just affect membrane diffusion, but could also alter interactions of the EnvCT with host cell proteins or underlying cytoskeleton machinery that are required for polarised recruitment of Env to the VS (Jolly *et al*, 2004). The increased diffusion of W757A Env provides a plausible explanation for decreased W757A Env at the VS and suggests that the polarisation of Env to sites of cell-cell contact is not mediated simply by immobilising surface-associated Env following binding to CD4 on target cells, but rather that cell-cell contact must specifically trigger or signal Env recruitment to the contact zone. Consistent with this notion, W757A was competent for Env-CD4 and co-receptor binding by neutralisation assay. We also found that the mobility of this mutant at sites of virus assembly was unperturbed when compared to WT Env trimers, suggesting that conformational changes in the baseplate structure or membrane association of LLP-2 does not impact Env incorporation and retention in the Gag lattice. This is consistent with our biochemical observations that W757A showed no incorporation defect, demonstrating that once Env associates with the Gag lattice it will be incorporated into viral particles. Of note, while our data indicate that the earliest events of VS formation and cell-cell assembly of HIV-1 require correct and regulated Env movement within the plasma membrane, they do not exclude that subsequent Env recruitment may be mediated by additional mechanisms described above.

Env is known to be a key driver of VS formation and cell-cell spread and inhibiting Env-receptor interactions abolishes VS formation (Jolly *et al*, 2004). We now show that the contribution of Env is more than mediating stable cell-cell interactions and providing adhesive interactions, rather the EnvCT appears to signal in some way to regulate the recruitment of Gag to the VS, thus to control the timing and location of virus assembly. Mechanistically, these observations may be explained by two possibilities. Firstly, they may allude to a sequence of assembly events in which the EnvCT in the PM initiates Gag lattice formation and budding following contact with a susceptible target cell (Hübner *et al*, 2009). In fact, the EnvCT YxxL mutant has been reported to affect the site of virus budding in polarised epithelial cells and T cells (Deschambeault *et al*, 1999; Lodge *et al*, 1994), supportive of an unappreciated role for the EnvCT in regulating viral assembly. A second possibility is that EnvCT at the neck of the budding virions is essential to complete the budding process (Buttler *et al*, 2018) and that mutating a key residue the EnvCT inhibits this; however, while this may explain the defect in cell-free viral budding, it does not explain the dramatic lack of Gag polarisation to the VS. The initial engagement of Env and CD4/coreceptor results in polarisation of the cellular cytoskeleton to the contact site and disrupting the actin cytoskeleton inhibits VS formation as well as plasma membrane virus assembly platforms (Rudnicka *et al*, 2009; Chen *et al*, 2007; Jolly *et al*, 2004, 2007b). Whether the underlying host cell cytoskeletal or associated cellular proteins link the EnvCT to polarised Gag lattice formation in some way (via undefined host factors) is an interesting possibility. It cannot be excluded that direct interactions between the EnvCT and Gag take place; however, while evidence for a direct interactions have been shown biochemically *in vitro* (Cosson, 1996; Alfadhli *et al*, 2019) they have not been convincingly demonstrated consistently, and we were unable replicate this during the course of this study. Clearly, further work is needed to determine the molecular interactions that allow the EnvCT to regulate Gag localisation during viral egress.

In addition to a lack of VS formation, cell-cell spread was also affected by a defect in infectivity of the W757A mutant virus. This was initially surprising, given that many other features of Env W757A remained unchanged, and unlike EnvCT truncation (CTΔ144) that also abrogate cell-cell spread (Durham & Chen, 2015; Murakami & Freed, 2000; Checkley *et al*, 2011), EnvCT W757A had no incorporation defect and did not display altered cell surface expression or recycling in T cell lines. Further analysis revealed that this infectivity defect was explained by the W757A virus showing a striking defect in fusion and thus viral entry, indicating that a single residue change in LLP-2 distal to the fusion peptide significantly impacted fusion domain structure and function. In support of our data, it has been reported that the LLP-2 helix influences fusogenicity and that viruses containing EnvCT truncations are less sensitive to T20 inhibition (Wyss *et al*, 2005; Kalia *et al*, 2003; Lu *et al*, 2008). Recent structural insights of the EnvCT have identified the role of the LLP-2 helix to act as a baseplate that supports the Env transmembrane domain (TMD) and stabilises the MPER region of Env, holding both in the correct structural antigenic state (Piai *et al*, 2020). It is highly plausible that W757 in LLP-2 is key to maintenance of the EnvCT baseplate (Supplementary Fig. 5). Mutation of the hydrophobic W757 may be sufficient to perturb the baseplate, conferring conformational changes via the TMD (Piai *et al*, 2020) on the MPER and Env fusion domains, destabilising the Env ectodomain leading to reduced Env fusogenicity. Thus, our data support with the notion that the EnvCT baseplate plays a putative role in regulating viral entry (Piai *et al*, 2020). Of note, retroviruses that contain shorter EnvCTs like MLV exploit a C-terminal R peptide that controls membrane fusion activity, deletion of which lowers Env fusogenicity (Kubo *et al*, 2007). Thus it appears a key function of the long lentiviral EnvCT and its membrane associated structure may be to regulate fusion, and in the case of HIV-1, via LLP-2 and specifically residue W757.

In conclusion, here we have identified a single tryptophan residue in HIV-1 EnvCT that potently regulates HIV-1 assembly, virion fusion and cell-cell spread between T cells. That this W757 residue is highly-conserved in pandemic HIV-1 group M and its SIVcpz ancestor suggests to a fundamental role for conservation of this reside in HIV-1 replication. This may prove a promising future therapeutic target to interfere with HIV-1 assembly and inhibit the dominant mode of HIV-1 dissemination.

## Materials and Methods

### Cells and viral constructs

Jurkat CE6.1 T cells were grown in RPMI 1640 medium (Thermo Fisher Scientific) supplemented with 10% fetal calf serum (FCS, Labtech) and 100 U/ml Penicillin-Streptomycin (Thermo Fisher Scientific). HEK 293T and HeLa TZM-bl cell lines were grown in DMEM medium (Thermo Fisher Scientific) supplemented with 10% FCS, 100 U/ml Penicillin-Streptomycin. Peripheral blood mononuclear cells (PBMC) were isolated from buffy coats from healthy donors using Ficoll Paque Plus (Sigma) according to manufacturer’s instructions. CD4+ T cells were isolated by negative selection using MojoSort Human CD4 T Cell Isolation kit (Biolegend). Primary CD4+ T cells were maintained in complete RPMI 1640 medium supplemented with 10 IU/ml IL-2 (Center for AIDS Reagents [CFAR], National Institute of Biological Standards and Controls, UK). Primary CD4+ T cells were activated for 4-5 days on T25 flasks coated with 5μg anti-CD3 antibody (clone OKT3, Biolegend) in the presence of 2μg/ml soluble anti-CD28 antibody (clone CD28.2, Biolegend). The CEM-A cell line was obtained through NIH AIDS Reagent Program, Division of AIDS, NIAID, NIH: CEM-A from Dr. Mark Wainberg and Dr. James McMahon, CEM-CL10 (Tremblay *et al*, 1989). Complete growth media for the CEM-A cell line was prepared by combining 10% fetal bovine serum (Corning; Corning, NY, USA), 2 mM L-glutamine (Corning), 1% hypoxanthine, thymidine (HT) solution (Corning), and Penicillin-Streptomycin (Corning) into Roswell Park Memorial Institute (RPMI) medium (Corning). HIV-1 NL4.3 (donated by Dr. Malcolm Martin, obtained from CFAR). Site directed mutagenesis was carried out on full length replication competent pNL4.3 by 15 amplification cycles using Pfu polymerase with the following primers; EnvCT W757A (GTGAACGGATCCTTAGCACTTATC<u>GCG</u>GACGATCTGCGGAGCCTGTG), W757Y (GTGAACGGATCCTTAGCACTTATC<u>TAC</u>GACGATCTGCGGAGCCTGTG), W757F (GTGAACGGATCCTTAGCACTTATC<u>TTC</u>GACGATCTGCGGAGCCTGTG), and MA Q62R (GACAAATACTGGGA<u>CGG</u>CTACAACCATCCCTTCAG), followed by Dpn1 (NEB) digest and sequence verification. Viral stocks were generated by transfecting 293T cells using Fugene 6 (Promega). After 48h, virus was harvested and infectivity assayed by serial titration on HeLa TZM-bl reporter cells using Bright-Glo (Promega).

### Antibodies

Antibodies used for flow cytometry: anti-HIV-1 p24-PE (KC57-RD1 Beckman Coulter); anti-HIV-1 Env clone PGT151 (gift from Laura McCoy, UCL); Secondary antibody: anti-Human IgG Cy5 (polyclonal Bethyl). Antibodies for western blotting: anti-HIV-1 gp120 rabbit antisera (donated by Dr S. Ranjibar and obtained from the CFAR); anti-HIV-1 gp41 Mab 246-D (donated by Dr S. Zoller-Pazner and Dr M. Gorny, obtained from the CFAR); and anti-HIV-1 Gag rabbit antisera (donated by DR G. Reid and obtained from the CFAR). Secondary antibodies: anti-Rabbit IgG (ab216773, Abcam), anti-Mouse IgG (ab216775, Abcam) and anti-Human IgG (926-32232, Licor). The BG18-QD625 antibody fab fragment probe was produced recombinantly for single particle tracking as previously described (Groves *et al*, 2020).

### HIV-1 infections and cell-cell spread assays

T cells were infected with 50,000 TCID50/million cells with VSV-G pseudotyped HIV-1 by gravity infection for 4 hours. For cell-cell spread assays HIV-1 infected T cells (donors) were analysed for Gag expression by flow cytometry 48h post-infection (p.i.). Uninfected T cells (targets) were labelled with 2.5 μM Cell Proliferation Dye eFluor 450 (Thermo Fisher Scientific) according to manufacturer’s instructions. HIV-1 infected Gag+ donors and dye-labelled targets were mixed in 1:1 ratio, incubated at 37°C for indicated time points and analysed by flow cytometry using a BD LSR Fortessa X-20 or Calibur cytometer. Data were analysed using FlowJo software.

### Virus release and infectivity assays

HIV-1 infected T cells were incubated for 24h and virus-culture supernatants were harvested. Total virus released into the supernatant was measured by qPCR to quantify the supernatant reverse transcriptase (RT) activity using SG-PERT assay (Pizzato *et al*, 2009) and RT activity was normalised to the number of Gag+ (HIV-1 infected) cells. Virion infectivity was determined by luciferase assay using HeLa TZM-bl reporter cells. To determine particle infectivity, RLU infectivity values were normalized to RT activity.

### Viral Fusion Assay

NL4.3 WT or W757A was co-transfected with BlaM-Vpr (Addgene) and pAdVAntage (Promega) into 293T cells to generate virions packaging BlaM-Vpr. Virus was concentrated by ultracentrifugation and quantified by SG-PERT. Next, 10^11^ mU RT of virus was used to infect target 10^6^ Jurkat cells for 4 hours by gravity infection and the BlaM-Vpr assay was performed as described (Cavrois *et al*, 2002).

### Immunofluorescence

Uninfected Jurkat cells (target cells) were labelled with 10 μM CellTracker Blue CMAC dye (Thermo Fisher Scientific) according to the manufacturer’s instructions. Infected Jurkat cells (donor cells) were mixed with target cells in a 1:1 ratio and incubated with (nonblocking) anti-Env antibody for 1 h at 37°C on poly-L-lysine–coated coverslips (Jolly *et al*, 2004). Cells were fixed with 4% paraformaldehyde, permeabilised with 0.1% Triton X-100 and stained with anti-Gag serum. Primary antibodies were detected with appropriate fluorescent secondary antibodies. Coverslips were mounted with ProLong Gold Antifade mounting solution (Thermo Fisher Scientific). Images were acquired on a DeltaVision Elite image restoration microscope (Applied Precision) with softWoRx 5.0 software. Envelope enrichment at the VS was measured by the increase in fluorescence intensity at the contact site over intensity at a distal membrane region, normalised to background intensity. Image processing and analysis was performed using FIJI software (Schindelin *et al*, 2012).

### Simultaneous Super-resolution and Single-Particle Tracking of HIV-1 Biogenesis

All imaging, processing and analysis was done per the optimised methodology of Groves et. Al., 2020 (Groves *et al*, 2020). Briefly, CEM-A T cells in complete RPMI medium (10% fetal bovine serum; 2 mM L-glutamine; 1% penicillin/streptomycin, and 1% hypoxanthine/thymidine; Corning, USA) were seeded on 25 mm No. 1.0 glass coverslips (Warner Instruments; Hamden, CT, USA) and infected with VSV-G pseudotyped virus packaged with the pSV-NL4-3 HIV-1 reference genome, encoding the previously described CA-Skylan-S nanobody probe, with deletions in the *gag-p6* late-domain, *pol, vif, vpr*, and *nef* genes (Groves *et al*, 2020). Forty-eight hours post-infection, cells were stained with BG18-QD625 probe and coverslips were transferred to specimen holders and mounted on a custom-built ring-TIRF microscope with a live-cell chamber maintaining 37° C and 5% CO_2_. Skylan-S and QD625 were simultaneously excited with a 473 nm laser and fluorescence was detected by separation of wavelengths using a dichroic beamsplitter housed in a W-View Gemini image splitter (Hamamatsu, Hamamatsu City, Japan). As previously described, images were focused onto two halves of a liquid cooled ORCA Fusion scientific-CMOS camera (C14440-20UP) streaming at 100 Hz (Groves *et al*, 2020). Raw images were corrected for non-uniform pixel offset and split based on wavelength. Single-molecule localisations for both channels were fit using custom software (IDL, Harris Geospatial; Broomfield, CO, USA), and lateral chromatic aberration between channels was corrected using a high density fiducial map collected after data acquisitions (100 nm TetraSpeck; Thermo Fisher Scientific). Single-molecule localisations were passed to an automated package written in MATLAB (Natick, MA). This software determined Gag assembly centroids and linked single molecule Env trajectories from frame to frame. Tracks were split into ‘proximal’ and ‘distal’ classifications based on the proximity of their localisations to Gag bud centroids as previously described (Groves *et al*, 2020); Linking parameters: 50 ms maximum time gap, 10 localisations per track minimum, 1 pixel [108.33 nm] maximum frame-to-frame distance). The apparent trajectory diffusion coefficients (D_apparent_) and the slope of the moment scaling spectrum (S_MSS_) were computed by fit to a linear regression and weighted by localisation uncertainties.

### Envelope Recycling Assay

Infected Jurkat cells were stained with Envelope specific antibodies PGT151 Ab (5 μg/mL), and 2G12 (5 μg/mL) for 60 min at 37°C in FCS supplemented RPMI to allow for Env internalisation (Mesner *et al*, 2020). Cells were thoroughly washed in ice cold FACS wash buffer and surface Env was detected with anti–human-PE secondary antibody for 30 min on ice to measure Env surface levels at T_0min._ Cells were again washed extensively in ice cold buffer and incubated with anti-human-Cy5 secondary antibody for 0 to 180 min at 37°C in FCS supplemented RPMI to label Env that is recycling back to the cell surface. Cells were fixed in 4% paraformaldehyde and analysed by flow cytometry.

### Flow cytometry

T cells were washed and incubated with antibodies (described above) at 4°C for surface staining and cell viability dye Zombie UV (Biolegend). Primary antibodies were detected with anti-human IgG secondary antibody, fixed and analysed. To stain for intracellular antigens, cells were fixed with 4% paraformaldhyde and permeabilised with CytoPerm buffer (Biolegend). Cells were analysed on BD LSR Fortessa X-20 cytometer. Compensation was performed using CompBeads (BD) and calculated by FACSDiva software. Data was analysed using FlowJo software.

### Western Blotting

Fifty μg of cell lysate and an equivalent volume of purified virus (purified by ultracentrifugation over sucrose) were separated by SDS-PAGE and analysed by western blotting using the following primary antibodies: anti-HIV-1 gp120, anti-HIV-1 gp41, anti-HIV-1 Gag, and anti-tubulin (described above). Primary antibodies were detected with appropriate fluorescent secondary antibodies and imaged with Odyssey Infrared Imager (Licor). Immunoblots were analysed with Image Studio Lite software.

### Neutralisation assays

Antibody and inhibitor neutralisation assay was performed as described previously (Montefiori, 2009). Briefly, virus supernatants were incubated with serial dilutions of soluble CD4 (CFAR NIBSC), AMD3100 (CFAR, NIBSC), PGT151 (Falkowska *et al*, 2014) (gift from Laura McCoy, UCL) or T-20 fusion inhibitor (CFAR, NIBSC) and incubated at 37°C for 1h. The mixture was added to TZM-bl cells and luciferase activity measured 48h later using Bright-Glo substrate. Neutralisation was calculated as percent decrease in luciferase activity compared to virus only control. IC_50_ values were calculated by non-linear regression analysis (sigmoid curve interpolation) using Prism software (GraphPad).

### Statistical analysis

Statistical significance was calculated where appropriate using a paired/unpaired student’s *t* test or one way ANOVA. Significance was assumed when p<0.05. All statistical analyses were calculated using Prism 9 (GraphPad Prism).

## Acknowledgments

This work was funded by a Wellcome Trust Investigator award (108079/Z/15/Z) to C.J. SvE is supported by a grant from the National Institute of Allergy and Infectious Diseases, R01AI138625. We are grateful to Ann-Kathrin Reuschl and Maitreyi Shivkumar for helpful discussions. We also acknowledge Laura McCoy (UCL), Petra Mlcochova (DoM, Cambridge), the NIBSC Centre for AIDS Reagents and the NIH AIDS Reagent Program for reagents.

## Figure Legends

Figure 1. EnvCT W757A does not polarise to viral synapses and leads to a defect in cell-cell spread. **(A)** Schematic showing the HIV-1 NL4.3 EnvCT sequence and location of W757 in the LLP2 alpha helix. Logo plots (generated using WebLogo3) indicate the probability of 5916 HIV-1 Env sequences analysed in the LANL database containing a W residue at the indicated positions. **(B)** Synapse formation between Jurkat cells infected with WT NL4.3 (left panel) or the W757A mutant (right panel) and uninfected target cells. NL4-3 virus pseudotyped with VSV-G to normalise viral entry was used to infect donor Jurkat cells. Representative image is an *xy* slice through the middle of a cell-cell contact between an infected donor cell (Env, green; Gag, red) and a target cell stained with a cell trace dye (blue). **(C)** Percent of contacts that exhibited synapse formation (defined by polarisation of both Env and Gag to the contact site). **(D)** Quantification of the percentage of infected donor cells (Gag+) in contact with target cells (blue). Data show the mean and SEM from n=54-69 infected donor cells compared using a two-tailed paired t test (ns, not significant; *** p < 0.001). **(E)** Signal intensity enrichment at the synapse, relative to a synapse distal site was determined from 20 images using ImageJ analysis. **(F)** Infected donor cells were mixed in a 1:1 ratio with target cells (stained with cell trace far red) and viral spread was measured by intracellular Gag staining in target cells by flow cytometry. Data show the percent of Gag+ target cells over time. Jurkat cells (left) and primary CD4+ cells (right). Data show the mean and SEM from four independent Jurkat cells experiments analysed using multiple paired t tests (ns, not significant; *** p < 0.001). For primary CD4^+^ T cells a representative donor from three unique donors is shown.

Figure 2. W757A EnvCT does not display a Gag lattice trapping defect at virus assembly sites. CEM-A adherent T cells were transduced with WT and W757A mutant Envelope virus and imaged after 48 hours. **(A)** Super-resolution reconstructions of Gag and lattice proximal/distal single-molecule localisations of Env are demonstrated (Scale bars = 200nm. Arrows highlight highly confined single molecule localisations of Env). Proximal and distal single-molecule localisations show the confined and freely-diffusing nature of single Env trimers, respectively. **(B)** Wild-type and mutant Env trimers do not differ in mobility when lattice proximal (top panels) (WT: *D*_*apparent*_ = 0.038 ± 0.06 μm^2^s^−1^, *S*_*MSS*_ = 0.03 ± 0.13; W757A: *D*_*apparent*_ = 0.033 ±0.03 *μ*^2^*s*^−1^, *S*_*MSS*_ =0.045 ±0.16), whereas lattice-distal diffusion differs significantly between viruses (bottom panels) (WT: *D*_*apparent*_ = 0.07 ± 0.17 *μ*^2^*s*^−1^, *S*_*MSS*_ = 0.12 ± 0.21; W757A: *D*_*apparent*_ = 0.05 ± 0.10 *μ*^2^*s*^−1^, *S*_*MSS*_ = 0.13 ± 0.22). Statistical significance was determined by unpaired two-sample t-test. 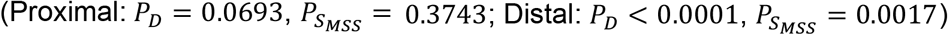. Error represents S.D.

Figure 3. EnvCT W757 is required for HIV-1 budding and infectivity. (A) Virus release from infected Jurkat and primary CD4^+^ T cells at 48hpi measured by quantifying RT activity in viral supernatants by SG-PERT qPCR. **(B)** Infectivity of virus particles released 48hpi was measured by titrating virus on HeLa TZM-bl reporter cells and RLU normalised per RT unit to calculate relative particle infectivity. **(C)** Western blot of cell lysate and purified virus from infected primary CD4^+^ T cells at 48hpi probed with anti-Env (gp120/gp160) and anti-Gag (p55/p24). **(D)** Quantification of Env (gp120) incorporation into virions normalised to Gag (p24) density, produced in Jurkat and primary CD4^+^ T cells. Data show the mean and SEM from at least three independent experiments compared using a two-tailed paired t test (ns, not significant; **** p < 0.0001).

Figure 4. Synthesis, processing, and recycling of EnvCT W757A is equivalent to WT EnvCT. **(A)** Intracellular staining of Env (green) and Gag (red) in Jurkat cells infected with WT and W757A at 48hpi. **(B)** Western blot of infected Jurkat cell lysates harvested 48hpi and probed with anti-gp120, anti-gp41, and anti-Gag p55/p24. Quantification of Env processing (gp120/gp160). Data show the mean and SEM from at least three independent experiments compared using a two-tailed paired t test (ns, not significant). **(C)** Flow cytometry analysis of cell surface Env levels on cells infected with WT (black) and W757A (grey) virus measured using bNAb PGT151. Uninfected control is shown in light grey. **(D)** Time course of endocytosed Env recycling back to the plasma membrane measured by flow cytometry. Env was stained with either 2G12 or PGT151. Envelope recycling was measured by staining with envelope antibody and a PE-secondary on ice followed by Cy5 at 37°C. Recycling is shown as the MFI of PE^+^CY5^+^ signal on the surface over time. Data show the mean and SEM from three independent experiments.

Figure 5. W757A virus shows a defect in viral fusion **(A)** Neutralisation of WT and W757A virus (produced from Jurkat T cells) by PGT151 bNAb, sCD4, AMD3100 and T20. IC_50_ values calculated by non-linear regression analysis of the neutralisation curves. ND denotes not determined. **(B)** W757A Env is defective in viral fusion compared to WT virus as measured by the BlaM-Vpr fusion assay. Data are the mean and SEM compared using an unpaired t test (*** p< 0.001). Mock refers to co-culture between uninfected T cells and target T cells.

Figure 6. Conservation of W757 across HIV and SIV lineages. **(A)** Amino acid alignment of the LLP2 domain of the EnvCT from different HIV and SIV consensus sequences. See also Supplementary Figure S4. **(B-D)** Viral budding, infectivity, and cell-cell spread can be rescued by substitution at position 757 with either tyrosine Y, or phenylalanine, F. Data are the mean and SEM compared using a two-tailed paired t test (ns, not significant; *** p< 0.001).

Supplementary Figure 1. EnvCT W757A is defective in T cell – T cell spread. Representative flow cytometry data of cell-cell spread between Jurkat T cells **(A)** and primary CD4^+^ T cells **(B)**. The percentage of cells in each quadrant is indicted.

Supplementary Figure 2. EnvCT W790A virus does not perturb HIV-1 Env function and viral cell-cell spread. **(A)** Alanine substitution of the tryptophan at HIV-1 envelope position 790 does not affect the ability of infected Jurkat cells to form conjugates with uninfected target cells. **(B-C)** W790A infected cells also form synapses to the same extent at WT HIV-1 and result in no defect in both intracellular and surface localization of Env. **(D)** Quantification of Env (gp120) incorporation into virions produced in Jurkat cells, normalised to Gag (p24) density. **(E)** Cell surface W790A Envelope (grey) measured by flow cytometry is equal to WT (black) levels. **(F-G)** Budding of virons from infected cells, and the relative infectivity of virions is equivalent to WT. (H) Viral transfer via cell-cell spread is the same with WT Env and W790A Env, but is significantly defective in W757A Env infection.

Supplementary Figure 3. Compensatory Matrix mutant (Q62R) is unable to alleviate the W757A phenotype. MA mutation Q62R does not rescue the defect in W757A budding **(A)** or infectivity **(B)**. Data are the mean and SEM compared using a one way ANOVA (ns, not significant; ** p< 0.01; **** p< 0.0001).

Supplementary Figure 4. Conservation of W757 among HIV-1 M group, SIVcpz, and SIVgor envelope glycoprotein. Phylogenetic analysis of the EnvCT showing amino acid residues found at position 757 (based on 194 consensus envelope sequences from http://www.hiv.lanl.gov/) performed using Chromaclade. Tryptophan (W) – dark blue. Tyrosine (Y) – light blue. Isoleucine (I) – purple. Leucine (L) – turquoise.

Supplementary Figure 5. W757 within the EnvCT LLP2 helix may stablise the EnvCT baseplate. (A) Top down and (B) side on models of trimeric EnvCT and TMD within the plasma membrane. Trytophan (red) and Alanine (blue) are highlighted in (C). The aromatic side chain of W757 (indicated by the yellow arrow) is predicted to be buried into the membrane, maintaining the EnvCT baseplate conformation, whereas alanine substitution at this position allows for more conformational flexibility in the quaternary structure of the EnvCT baseplate, disrupting the TMD and perhaps MPER/gp120 conformations.

---

**Supplementary Figure 1.**
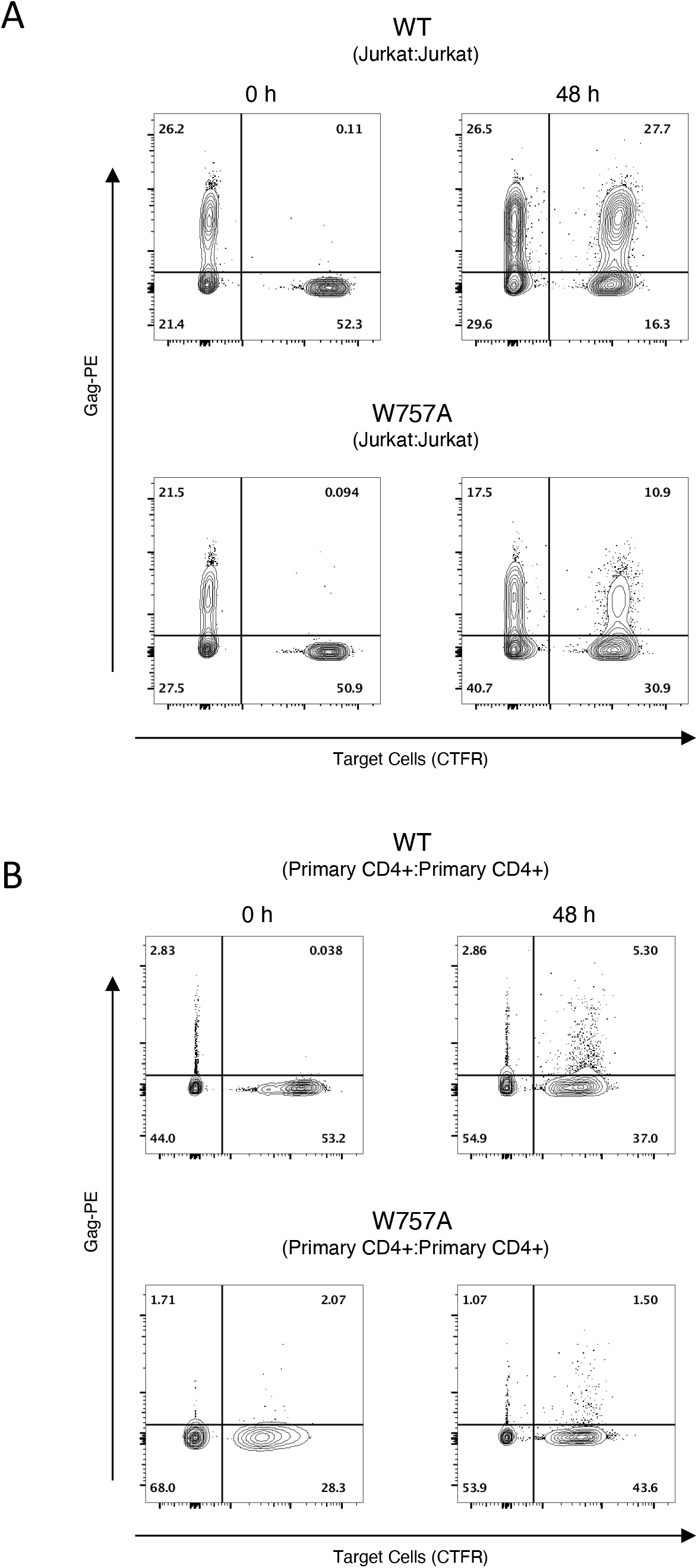
EnvCT W757A is defective in T cell – T cell spread. Representative flow cytometry data of cell-cell spread between Jurkat T cells **(A)** and primary CD4^+^ T cells **(B)**. The percentage of cells in each quadrant is indicted.

**Supplementary Figure 2.**
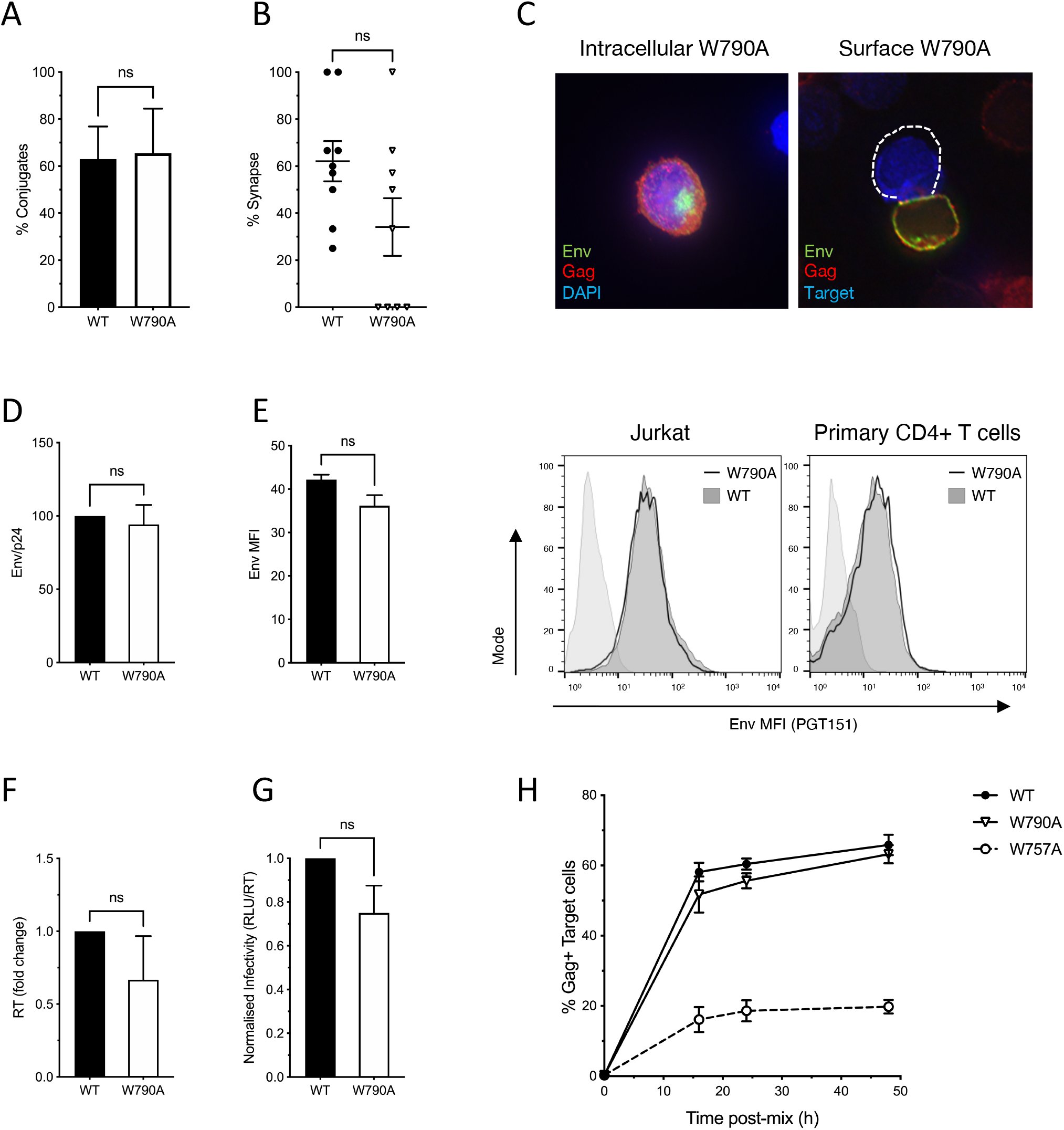
EnvCT W790A virus does not perturb HIV-1 Env function and viral cell-cell spread. (A) Alanine substitution of the tryptophan at HIV-1 envelope position 790 does not affect the ability of infected Jurkat cells to form conjugates with uninfected target cells. **(B-C)** W790A infected cells also form synapses to the same extent at WT HIV-1 and result in no defect in both intracellular and surface localization of Env. **(D)** Quantification of Env (gp120) incorporation into virions produced in Jurkat cells, normalised to Gag (p24) density. **(E)** Cell surface W790A Envelope (grey) measured by flow cytometry is equal to WT (black) levels. **(F-G)** Budding of virons from infected cells, and the relative infectivity of virions is equivalent to WT. **(H)** Viral transfer via cell-cell spread is the same with WT Env and W790A Env, but is significantly defective in W757A Env infection.

**Supplementary Figure 3.**
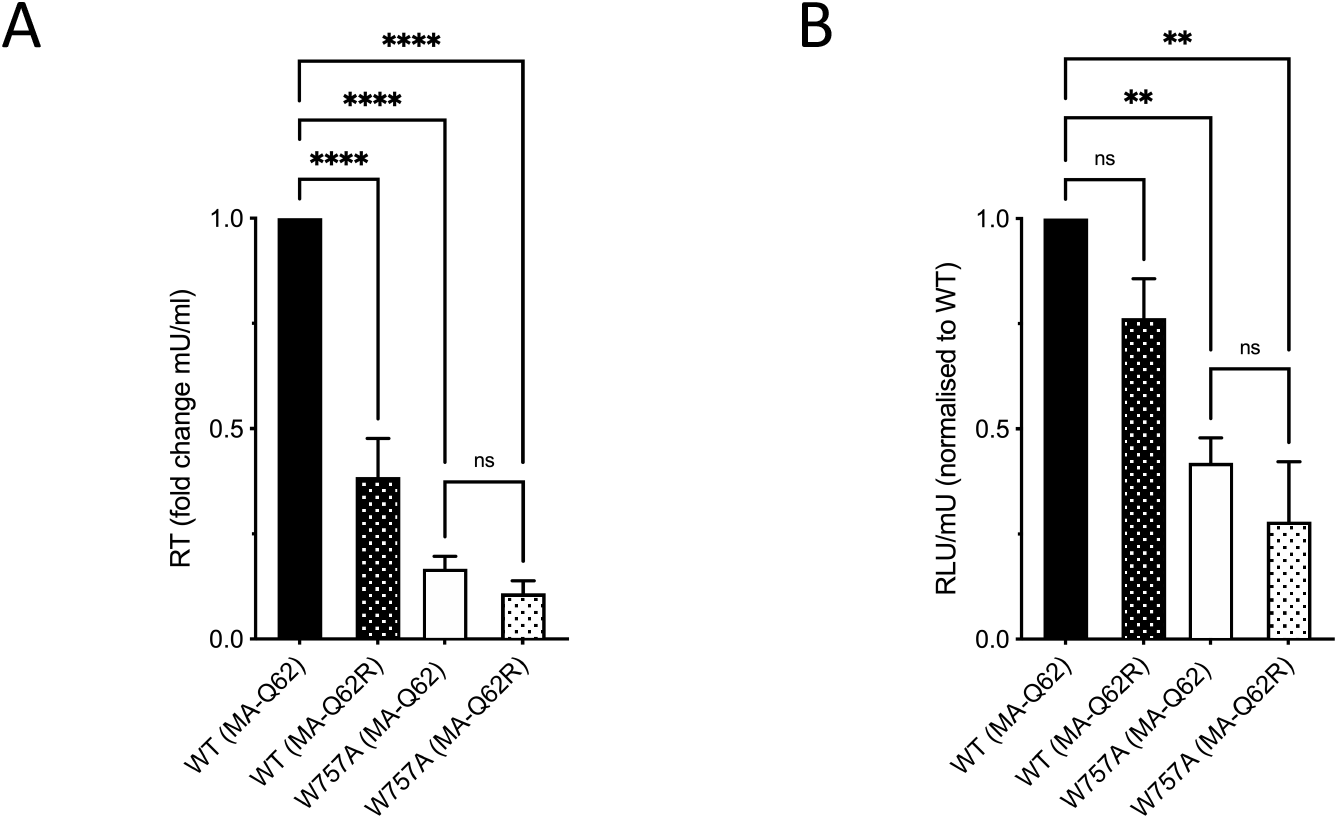
Compensatory Matrix mutant (Q62R) is unable to rescue W757A. MA mutation Q62R does not rescue the defect in W757A budding **(A)** or infectivity **(B)**. Data are the mean and SEM compared using a one way ANOVA (ns, not significant; ** p< 0.01; **** p< 0.0001).

**Supplementary Figure 4.**
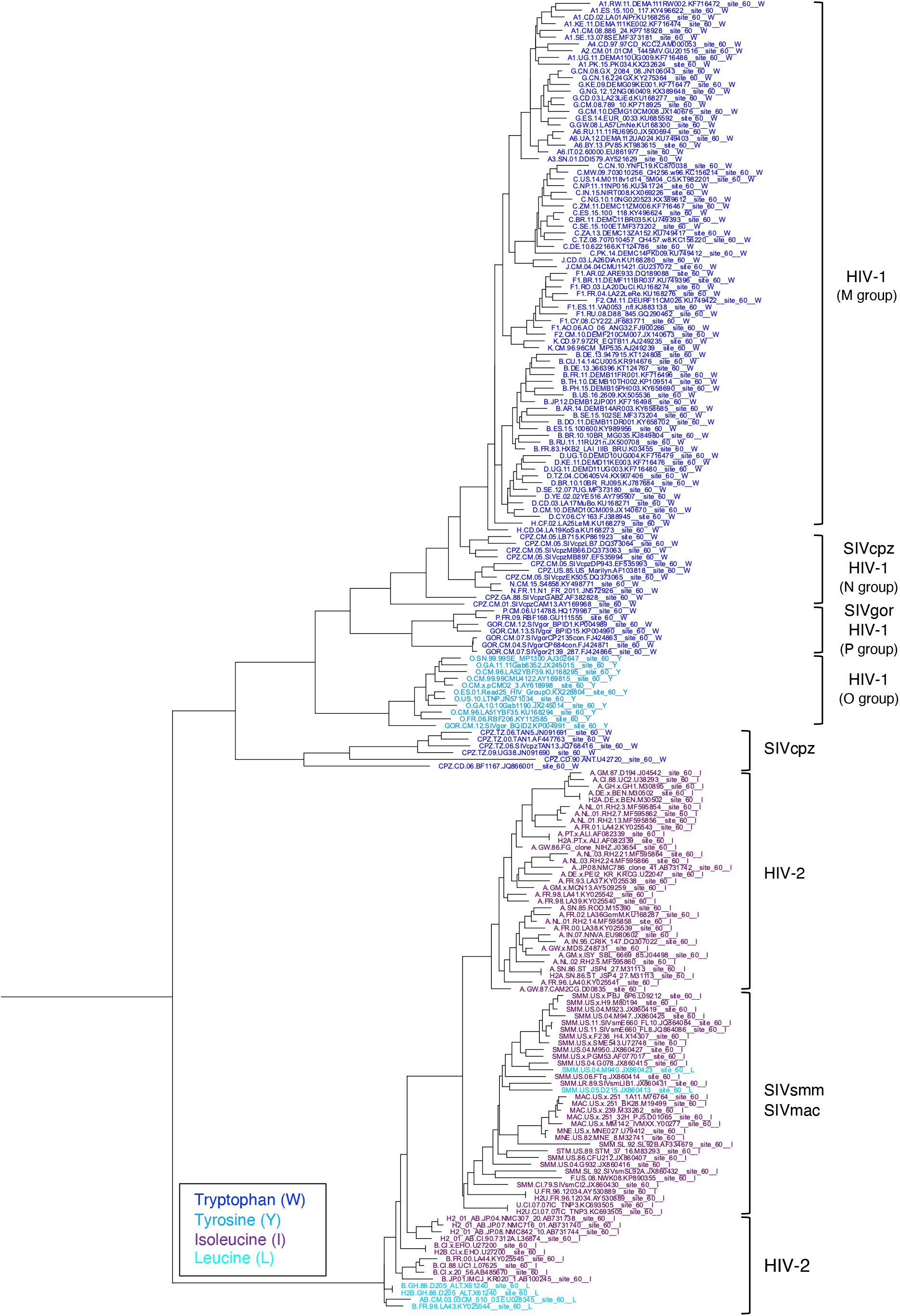
Conservation of W757 among HIV-1 M group, SIVcpz, and SIVgor envelope glycoproteins. Phylogenetic analysis of the EnvCT showing amino acid residues found at position 757 (based on 194 consensus envelope sequences from http://www.hiv.lanl.gov/) performed using Chromaclade. Tryptophan (W) – dark blue. Tyrosine (Y) – light blue. Isoleucine (I) – purple. Leucine (L) – turquoise.

**Supplementary Figure 5.**
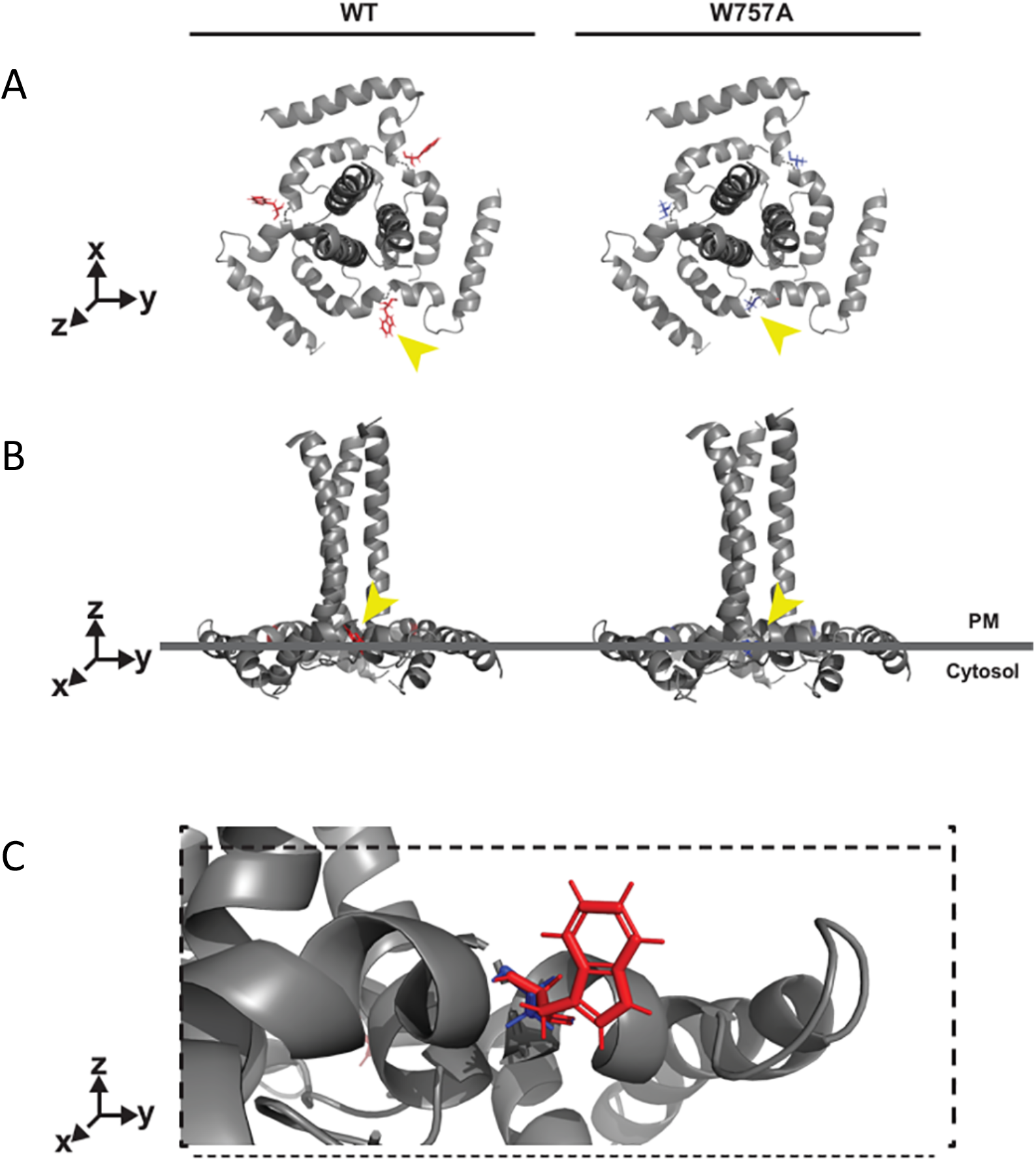
W757 within the EnvCT LLP2 helix may stablise the EnvCT baseplate. (A) Top down and **(B)** side on models of trimeric EnvCT and TMD within the plasma membrane. Trytophan (red) and Alanine (blue) are highlighted in **(C)**. The aromatic side chain of W757 (indicated by the yellow arrow) is predicted to be buried into the membrane, maintaining the EnvCT baseplate conformation, whereas alanine substitution at this position allows for more conformational flexibility in the quaternary structure of the EnvCT baseplate, disrupting the TMD and perhaps MPER/gp120 conformations.

